# A Novel Ensemble-Based Machine Learning Algorithm To Predict The Conversion From Mild Cognitive Impairment To Alzheimer’s Disease Using Socio-demographic Characteristics, Clinical Information And Neuropsychological Measures

**DOI:** 10.1101/564716

**Authors:** Massimiliano Grassi, Nadine Rouleaux, Daniela Caldirola, David Loewenstein, Koen Schruers, Giampaolo Perna, Michel Dumontier, for the Alzheimer’s Disease Neuroimaging Initiative

**Author notes:** corresponding author Massimiliano Grassi. Co-first authors (Massimiliano Grassi, Nadine Rouleaux). **Authors’ email addresses** Nadine Rouleaux, Daniela Caldirola, David Loewenstein, Koen Schruers, Giampaolo Perna, Michel Dumontier. Data used in preparation of this article were obtained from the Alzheimer’s Disease Neuroimaging Initiative (ADNI) database (adni.loni.usc.edu). As such, the investigators within the ADNI contributed to the design and implementation of ADNI and/or provided data but did not participate in analysis or writing of this report. A complete listing of ADNI investigators can be found at http://adni.loni.usc.edu/wp-content/themes/freshnews-dev-v2/documents/policy/ADNI_Acknowledgement_List%205-29-18.pdf.

## Abstract

**Background:** Despite the increasing availability in brain health related data, clinically translatable methods to predict the conversion from Mild Cognitive Impairment (MCI) to Alzheimer’s disease (AD) are still lacking. Although MCI typically precedes AD, only a fraction of 20-40% of MCI individuals will progress to dementia within 3 years following the initial diagnosis. As currently available and emerging therapies likely have the greatest impact when provided at the earliest disease stage, the prompt identification of subjects at high risk for conversion to full AD is of great importance in the fight against this disease. In this work, we propose a highly predictive machine learning algorithm, based only on non-invasively and easily in-the-clinic collectable predictors, to identify MCI subjects at risk for conversion to full AD.

**Methods:** The algorithm was developed using the open dataset from the Alzheimer’s Disease Neuroimaging Initiative (ADNI), employing a sample of 550 MCI subjects whose diagnostic follow-up is available for at least 3 years after the baseline assessment. A restricted set of information regarding sociodemographic and clinical characteristics, neuropsychological test scores was used as predictors and several different supervised machine learning algorithms were developed and ensembled in final algorithm. A site-independent stratified train/test split protocol was used to provide an estimate of the generalized performance of the algorithm.

**Results:** The final algorithm demonstrated an AUROC of 0.88, sensitivity of 77.7%, and a specificity of 79.9% on excluded test data. The specificity of the algorithm was 40.2% for 100% sensitivity.

**Discussion:** The algorithm we developed achieved sound and high prognostic performance to predict AD conversion using easily clinically derived information that makes the algorithm easy to be translated into practice. This indicates beneficial application to improve recruitment in clinical trials and to more selectively prescribe new and newly emerging early interventions to high AD risk patients.

## Background

Alzheimer’s Disease (AD), the most common form of dementia, is a neurodegenerative disease characterized by progressive memory loss, cognitive impairment and general disability. The progression of AD comprises a long, unnoticed preclinical stage, followed by a prodromal stage of Mild Cognitive Impairment (MCI) that leads to severe dementia and eventually death [1]. While no disease-modifying treatment is currently available for AD, a large number of drugs are in development and encouraging early-stage results from clinical trials provide for the first time a concrete hope that one or more therapies may become available in a few years [2]. As the progression of the neuropathology in AD starts years in advance before clinical symptoms of the disease become apparent and progressive neurodegeneration has irreversibly damaged the brain, emerging treatments will likely have the greatest effect when provided at the earliest disease stages. Thus, the prompt identification of subjects at high risk for conversion to AD is of great importance.

The ability to identify declining individuals at the prodromal AD stage provides a critical time window for early clinical management, treatment & care planning and design of clinical drug trials [3]. Precise identification and early treatment of at risk subjects would stand to improve outcomes of clinical trials and reduce healthcare costs in clinical practice. However, simulations also suggest that the health care system is not prepared to handle the potentially high volume of patients who would be eligible for treatment [2].

MCI represents (currently) the earliest clinically detectable stage of progression towards AD or other dementias. To determine whether MCI symptoms are due to an AD pathology requires the use of blood testing, brain imaging and cerebrospinal fluid (CSF) biomarkers, in addition to careful medical evaluation of medical history, neuropsychological testing, and a physical and neurological examination. The cognitive decline in MCI is abnormal given an individual’s age and education level, but does not interfere with daily activities, and thus does not meet criteria for AD. However, only 20-40% of individuals will progress to AD within three years, with a lower rate of conversion reported in epidemiologic samples than in clinical ones [4,5]

Currently, there are no means to provide patients diagnosed with MCI with an early prognosis for conversion to full AD. While changes in several biomarkers prior to developing AD have been reported, no single biomarker appears to adequately predict the conversion from MCI to AD with an acceptable level of accuracy. As such, there is increasing evidence that the use of a combination of biomarkers can best predict the conversion to AD [3,6–9].

In the current age of big data and artificial intelligence technologies, considerable effort has been dedicated in developing machine learning algorithms that can predict the conversion to AD in subjects with MCI using different combinations of AD biomarkers with varying accuracy [7,10–18] (see [19,20] for a recent review of the most performing algorithms presented in the scientific literature so far). Many studies focused on predicting the conversion of AD in MCI patients, combine different types of data including brain imaging, CSF biomarkers, genotyping, demographic and clinical information, and cognitive performance.

However, while combining different biomarkers improves model accuracy, there is a lack of consistency regarding a specific combined AD prediction model and translation into practice is still lacking. One possible reason for this is that current algorithms generally rely on expensive and/or invasive predictors, such as brain imaging or CSF biomarkers. As such, these studies only serve the purpose of a proof-of-concept, without being further tested in independent and clinical samples.

The current study aimed to develop a clinically translatable machine learning algorithm to predict the conversion to full AD in subjects with MCI within a 3-year period, based on fast, easy, and cost-effective predictors. Our hypothesis was that high predictive accuracy could be obtained using a machine learning model with simple and non-invasive predictors. We used data obtained from the Alzheimer’s Disease Neuroimaging Initiative (http://adni.loni.usc.edu/) with a particular consideration for socio-demographic and clinical information, and neuropsychological test scores rather than using complex, invasive, and expensive imaging or CSF predictors.

## Materials and methods

### ADNI

Data used in the preparation of this article were obtained from the Alzheimer’s Disease Neuroimaging Initiative (ADNI) database (adni.loni.usc.edu). The ADNI was launched in 2003 as a public-private partnership, led by Principal Investigator Michael W. Weiner, MD. The primary goal of ADNI has been to test whether serial magnetic resonance imaging (MRI), positron emission tomography (PET), other biological markers, and clinical and neuropsychological assessment can be combined to measure the progression of MCI and early AD. It contains data of a large number of cognitive normal, MCI, and AD subjects recruited in over 50 different centers in US and Canada with follow-up assessments performed every 6 months.

For this study, we used a subset of the ADNI dataset called ADNIMERGE that includes a reduced selection of more commonly used variables (i.e. demographic, clinical exam total scores, MRI and PET variables). This subset is part of the official dataset provided by ADNI.

### Subjects

Data regarding 550 subjects with MCI and with available diagnostic follow-up assessments for at least three years were included in the study. The most relevant inclusion criteria of ADNI studies are the following: age between 55-90; six grade education or work history; subjects had to be fluent English/Spanish speakers; Geriatric Depression Scale score less than 6; good general health; no use of excluded medications (e.g. medications with anticholinergic properties) and stability for at least 4 weeks of other allowed medications; Hachinski ischemic score scale less than or equal to 4. A complete description of the ADNI study inclusion/exclusion criteria, including the full list of excluded and permitted medications, can be found in the ADNI General Procedure Manual, pages 20-25 (link: https://adni.loni.usc.edu/wp-content/uploads/2010/09/ADNI_GeneralProceduresManual.pdf). The diagnosis of MCI was performed with the following criteria: memory complaint by subject or study partner that is verified by a study partner; abnormal memory function documented by scoring below the education adjusted cutoff on the Logical Memory II subscale (Delayed Paragraph Recall) from the Wechsler Memory Scale – Revised, which is less than or equal to 11 for 16 or more years of education, less than or equal to 9 for 8-15 years of education, and less than or equal to 6 for 0-7 years of education; Mini-Mental State Exam (MMSE) score between 24 and 30; Clinical Dementia Rating (CDR) score of 0.5; Memory Box score at least of 0.5; general cognition and functional performance sufficiently preserved such that a diagnosis of AD cannot be made.

Subjects were classified as converters to probable AD (cAD; n = 197, 35.82%) if they satisfied the National Institute of Neurological and Communicative Disorders and Stroke/Alzheimer’s Disease and Related Disorders Association criteria for AD [28] during at least one of the follow-up assessments occurred within three years from the baseline investigation, as well as having a MMSE score between 20 and 2. Otherwise, they were classified as non-converters to AD (NC; n = 353, 64.18%).

### Feature extraction

Considering our aim to employ only predictors that are either already routinely assessed or easily introducible in clinical practice, and that are not perceived as invasive by patients, we decided to take into account only variables in the ADNIMERGE dataset that regards diagnostic subtypes, sociodemographic characteristics, clinical and neuropsychological test scores. Some of these variables were not available for all recruited subjects and it was a priori decided to remove variables with greater than 20% missing values. Only the Digit Span Test score (DIGIT) exceeded the cut-off (52.73%) and was not used in our analysis. The following variables were used:

- *Sociodemographic characteristics*: gender, age (in years), years of education, and marital status (never married, married, divorced, widowed, unknown).
- *Subtypes of MCI*: Early or Late MCI according to their score in the Logic Memory subscale of the Wechsler Memory Scale - Revised [21], adjusted for the years of education. 9-11 Early MCI and ≤8 Late MCI for 16 or more years of education; 5-9 Early MCI and ≤4 Late MCI for 8-15 years of education; 3-6 Early MCI and ≤2 Late MCI for 0-7 years of education.
- *Clinical scales*: CDR [22] was used to characterize six domains of cognitive and functional performance in AD and related dementias: Memory, Orientation, Judgment & Problem Solving, Community Affairs, Home & Hobbies, and Personal Care. The rating is obtained through a semi-structured interview of the patient together with other informants (e.g., family members). Sum of Boxes score was used in the current analyses (CDRSB). The score of the Functional Assessment Questionnaire (FAQ) [23], an a informant-based clinician-administered questionnaire which assess the functional daily-living impairment in dementia, was also used in the analyses.
- *Neuropsychological tests*: MMSE [24] is a 30-point questionnaire that is used measuring cognitive impairment. All MCI subjects has a score of 24 of more at baseline. The Cognitive Subscale Alzheimer’s Disease Assessment Scale (ADAS) [25] is made of 11 tasks that include both subject-completed tests and observer-based assessments, assessing the memory, language, and praxis domains. The result is a global final score ranging from 0 to 70, based on the sum of the scores of the single tasks (ADAS11). Beyond the ADAS11 score, the ADNI study included also an additional test of delayed word recall and a number cancellation or maze task, which are further summed to have a new total score that ranges from 0 to 85 (ADAS13). In addition, the score of the task 4 (Word Recognition, ADASQ4) was included in the ADNIMERGE dataset. All these three ADAS scores were initially considered as predictors in the analyses. The Rey Auditory Verbal Learning Test (RAVLT) [26] is a cognitive test used to evaluate verbal learning and memory. All the immediate (RAVLT-I), learning (RAVLT-L), forgetting (RAVLT-F), and percent forgetting (RAVLT-PF) scores were included in the ADNIMERGE dataset and used in the analyses. Moreover, the total delayed recall score of the Logic Memory subtest of the of the Wechsler Memory Scale-Revised [27] (LDT), which assess verbal memory, and the time to complete of the Trial Making Test version B (TMTBT) [28], which assess visual-motor coordination and attentive functions.

Taken together, 14 continuous, 2 dichotomous and 1 polytomous categorical features were initially considered. The full list is available in Table 1.

**Table 1.** Descriptive statistics.

| Continuous Predictors | Non-converters |  | Converters |  | Missing Values |  |
| --- | --- | --- | --- | --- | --- | --- |
|  | Mean | S.D. | Mean | S.D. | N | % |
| Age | 72.42 | 7.54 | 74.19 | 6.88 | / | / |
| Years of education | 16.18 | 2.74 | 15.74 | 2.83 | / | / |
| CDRSB | 1.26 | 0.70 | 1.95 | 1.01 | / | / |
| ADAS11 | 8.67 | 3.78 | 12.94 | 4.26 | 1 | 0.18% |
| ADAS13 | 13.89 | 5.81 | 21.05 | 5.72 | 3 | 0.55% |
| ADASQ4 | 4.61 | 2.35 | 7.16 | 2.04 | / | / |
| MMSE | 28.01 | 1.71 | 26.85 | 1.72 | / | / |
| RAVLT-I | 37.84 | 10.47 | 28.05 | 6.74 | / | / |
| RAVLT-L | 4.76 | 2.59 | 2.90 | 2.11 | / | / |
| RAVLT-F | 4.37 | 2.46 | 5.20 | 2.30 | / | / |
| RAVLT-PF | 51.09 | 30.92 | 78.20 | 28.04 | / | / |
| LDT | 6.84 | 3.12 | 3.59 | 2.89 | / | / |
| DIGIT | 40.24 | 10.42 | 34.86 | 11.02 | 290 | 52.73% |
| TMTBT | 100.30 | 49.56 | 141.24 | 79.66 | 4 | 0.73% |
| FAQ | 1.76 | 2.75 | 5.81 | 5.00 | 4 | 0.73% |

| Categorical Predictors |  | Non-converters |  | Converters |  | Missing Values |  |
| --- | --- | --- | --- | --- | --- | --- | --- |
|  |  | N | % | N | % | N | % |
| Gender | Male | 220 | 62.32% | 118 | 59.90% | / | / |
|  | Female | 133 | 37.68% | 79 | 40.10% |  |  |
| Subtype of MCI | Early | 22 | 6.23% | 22 | 11.17% | / | / |
|  | Late | 175 | 49.58% | 175 | 88.83% |  |  |
| Marital Status | Never Married | 6 | 1.70% | 3 | 1.52% | 3 | 0.55% |
|  | Married | 267 | 75.64% | 161 | 81.73% |  |  |
|  | Divorced | 35 | 9.92% | 13 | 6.60% |  |  |
|  | Widowed | 42 | 11.90% | 20 | 10.15% |  |  |

*S.D = Standard Deviation; N = numbers of subjects.*

### Dataset division in 5 site-independent, stratified test subsets

The entire dataset was divided in five mutually exclusive data subsets. These five subsets were created in order to satisfy the following criteria: every subset has to include roughly 20% of the cases; all subjects from each of the 58 different recruitment sites has to be allocated into the same subset; every subset has to include roughly the same percentage of cAD as observed in the entire dataset (35.82%). In order to accomplish a division in 5 folds which satisfies all these criteria, 10000 different subsets were generated by progressively adding all subjects from a randomly chosen recruiting site, until the included cases ranged between 19% and 21% of the entire sample. Then, only those subsets whose percentage of cAD ranged between 35.52% and 36.12% were retained, which was satisfied in 567 (5,67%) out of the generated subsets. Finally, all possible combinations of five of the retained subsets were created in order to identify whether in any of these combinations covered the entire dataset without any repetition of cases. The entire process took around 4 hours of computation (on a Linux server with 2.20GHz Intel Xeon E5-2650 v4 CPUs), and successfully found a single combination of five subsets that satisfied all the desired criteria (Table 2).

**Table 2.**
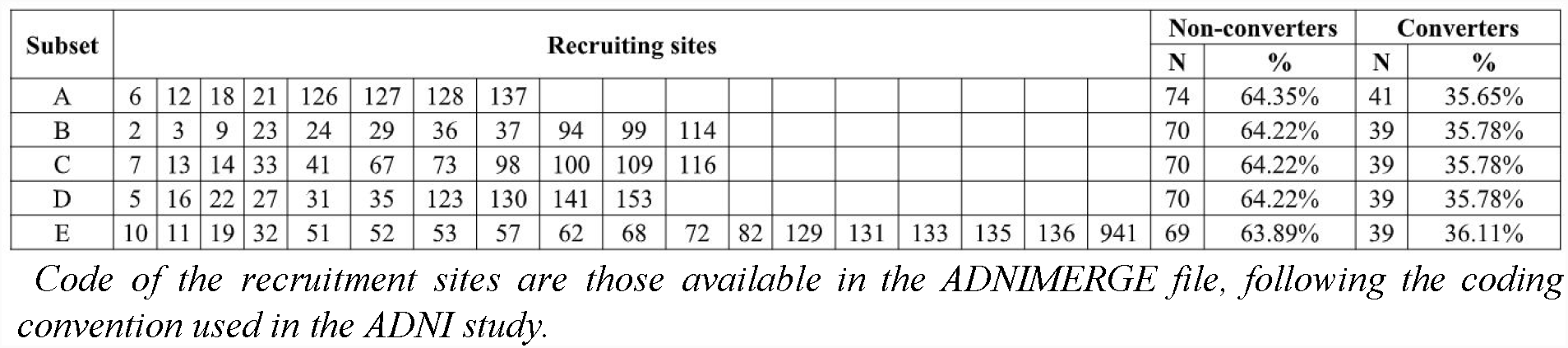
Allocation of the ADNI study recruitment sites in the five subsets.

All the missing value imputation, feature transformation and selection procedures, model training with cross-validation, and ensembling of different algorithms predictions described in the following paragraphs were performed in five distinct repetitions (named A-E) of the analyses, each time using the cases included in four of the five subsets and blindly to the remaining subset that were used as a test subset. The same missing value imputation, feature transformation and selection applied during training in the other four subsets were applied to the test subset. The predictive algorithms and their ensembling procedure developed in the other 4 subsets were tested against the test subset to obtain an estimate of the generalized performance in an independent sample of cases recruited in sites different from the ones used for training.

### Feature transformation and selection

Imputation was performed for variables with missing values using the median for continuous features and using the mode for categorical features. Continuous variables were standardized (mean = 0, standard deviation = 1) and non-dichotomous categorical variables were dichotomized using one-hot encoding, i.e. re-coding them in a new dichotomous variable for each class of the categorical variable, with 1 indicating the occurrence of that class and 0 the occurrence of any other class of the variable.

In case groups of variables resulted highly correlated (pairwise r >= .75), principal component analysis was used to calculate principal components and the original variables were substituted with all the components with eigenvalues >= 1.

All features were initially used during training (feature set 1). Moreover, three feature subsets were additionally created based on different selection strategies in order to include only those that are the most informative. A filtering procedure was applied to create reduced sets of features based on their bivariate statistical association (p < .05) with the outcome using independent sample t-test for continuous predictors and Fisher’s exact test for both dichotomous and one-hot encoded polytomous features (feature subset 2). Two cross-validated recursive feature elimination procedures (also known as “wrapper” procedures) with Logistic Regression (LR, feature subset 3) and Random Forest (RF, feature subset 4) [29] were also applied. In particular, the latter strategy was chosen because it has previously proved to be efficacious in selecting a relevant feature subset [19].

### Machine learning techniques

Several machine learning procedures that can be used to solve classification problems exists. We used 13 supervised techniques: LR, Naive Bayes (NB) [30], L1 and L2 regularized logistic regression or Elastic Net (EN) [31], Support Vector Machine [32] with linear (SVM-Linear), radial basis function (SVM-RBF), and polynomial (SVM-Poly) kernels with Platt scaling [33], k-Nearest Neighbors algorithm (kNN) [34], Multi-Layer Perceptrons with either one or two hidden layers and trained with either a full-batch gradient descent or adam [35] algorithms (MLP1-Batch, MLP2-Batch, MLP1-Adam, MLP2-Adam), RF, and Gradient Tree Boosting of Decision Trees (GTB) [36]. All analyses were parallelized on a Linux server equipped with four 12-core Intel Xeon CPU E5-2650 v4 @ 2.20GHz and were performed in Python 3.6 [37], using the implementation of the machine learning techniques available in the Scikit-Learn library [38].

### Hyper-parameter optimization

Machine learning techniques usually have one or more hyper-parameters that allow a different tuning of the algorithm during the training process. Different values of these hyper-parameters lead to algorithms with different predictive performances with the goal of obtaining the best possible performance when applied to cases that are not part of the training set. In order to optimize such hyper-parameters for each ML techniques used in this study, each model was trained with 50 random hyper-parameter configurations, and 50 further configurations were progressively estimated with a Bayesian optimization approach. Instead of a random generation, Bayesian optimization aims to estimate which is the hyper-parameter configuration that would maximize the performance of the algorithm starting from the previously attempted ones, based on the assumption that it exists a relationship between the various hyper-parameter values and the performance achieved by the algorithm. Bayesian optimization is expected to be able to identify better hyper-parameter configurations, and in a reduced number of attempts, compared to trying to generate them at random. Estimation was performed with Gaussian Processes, as implemented in the Scikit-Optimized library (https://scikit-optimize.github.io/).

The Area Under the Receiving Operating Curve (AUROC) was used as performance metric to be maximized. All the ML algorithms developed in this study output a continuous prediction score (range: 0-1; the closer to 1 the higher the predicted risk of conversion for that subject) and the AUROC value can be interpreted as the probability that a randomly selected cAD subject will receive a higher output score than a randomly selected NC subject. The AUROC value is 0.5 when the algorithm makes random predictions and 1 in case it is always correct in making predictions. AUROC is not affected by class imbalance and it is independent with respect to any specific threshold that is applied to perform a dichotomous prediction.

### Cross-validation procedure

The aim is to develop an algorithm that can achieve the best possible generalized performance and not to perform well only with the cases used in the training process. Cross-validation provides an estimate of such generalized performance for every hyper-parameter configuration. In cross-validation, the train sample is divided in several folds of cases that are held-out from the training process, with training iteratively performed with the remaining cases. After the training, the algorithm is finally applied on the held-out cases.

We applied the commonly used 10-fold cross-validation procedure, repeated 10 times to obtain a stable performance estimate. The fold creation was performed at random, stratifying (i.e. balancing) for the percentage of converters and non-converters in each fold. Finally, the 100 performance estimates of the algorithm available for each hyper-parameter configuration were averaged to provide a final point estimate of the generalized performance. The hyper-parameter configuration for each machine learning technique that demonstrated the best average cross-validated AUROC was retained.

### Weighted rank average of single algorithm predictions

Using a collection of algorithms and combining their predictions instead of considering only the prediction coming from a single algorithm generally improves the overall predictive performance [39]. This procedure is called ensembling and it is also the principles on which some individual techniques such as Random Forest and Gradient Boosting techniques are based.

Several different ensemble methods exist, which usually require a further independent data subset from both the training and test ones. This additional subset would be used to train how to optimally combine the various predictions generated by the single algorithms. Given the limited amount of data available in the current study, further reducing the size of the train sample may have undermined the predictive performance of the developed algorithms. Thus, we decided to apply a simple form of ensembling based on a weighted average of the rank predictions generated by all individual algorithms. This strategy is usually considered effective even though it does not require to develop any further machine learning meta-algorithm and to optimize its hyperparameters [40].

First, the ranks of the cross-validated continuous prediction scores of the train subset cases were calculated for each of the 52 developed algorithms, and rescaled in order to range between 0 and 1. Then, the arithmetic average of the rescaled ranks weighted for the cross-validated AUROC was calculated for each train subset case, representing the new continuous prediction scores for the train subset cases.

To generate the final continuous prediction scores of the test subset cases, at first 52 prediction scores for each test case were generated using all the 52 developed algorithms. Then, the prediction score of each algorithm was substituted with the rescaled rank of the closest cross-validated train subset prediction score of that algorithm. Finally, the average of the rescaled ranks weighted for the cross-validated AUROC was calculated. This represents the final continuous prediction scores of each test subset cases.

### Testing performance

The final continuous prediction scores of the five test subsets, which were obtained using the weighted rank average, were pooled and used to calculate the whole sample test AUROC. This represents the final estimate of the generalized site-independent AUROC that the algorithm is expected to achieve when it is applied to new cases. The 95% confidence interval (CI) of the AUROC was calculated with a stratified bootstrap procedure, with 10000 resamples and applying the bias-corrected and accelerated (BCa) approach [41].

Different categorical cAD/NC predictions were generated for each case applying various thresholds to the final continuous prediction scores (i.e. a score equal or above the threshold indicated a cAD, otherwise a NC). First, the threshold values that maximized the balanced accuracy (i.e. the average between sensitivity and specificity) of the cross-validated train subsample ensemble predictions in each of the five analyses replication was identified and averaged in order to have a final unique threshold that was applied to the final continuous prediction scores. Moreover, the threshold values that generated sensitivity of 100%, 97.5%, 95%, 90%, 85%, 80%, 75% of the cross-validated train subsample ensemble predictions in each of the five analyses replication was identified, averaged and applied to the final continuous prediction scores.

Specificity (i.e. recall), sensitivity, positive predictive value (i.e. precision), negative predictive value, balanced accuracy and F1 score (i.e. the harmonic average of the sensitivity and positive predictive value) were calculated considering the pooled categorical predictions generated with the abovementioned thresholds, which represent the estimates of the generalized site-independent performance of the algorithm when applied to perform categorical predictions of cAD/NC in new cases, such that either the balanced-accuracy is aimed to be maximized or defined levels of sensitivity are aimed to be obtained.

### Feature importance

To provide a general ranking of the importance of the predictors used in this study, we applied the same five train/test split protocol to iteratively develop logistic regression models using only a single feature, in the train subsets, and these models were applied to generate the continuous prediction scores in the five test subsamples. The scores of the test subsamples were finally pooled together and used to calculate the whole sample test AUROC for each predictor. This gives a metric of importance for each predictor that is independent from both the machine learning technique used and all other predictors inserted in the algorithm. The 95% confidence interval (CI) of also these AUROCs was calculated with a stratified bootstrap procedure, with 10000 resamples and applying the bias-corrected and accelerated (BCa) approach [41].

## Results

Descriptive statistics of each feature in the cAD and NC groups are reported in Table 1. Statistics of continuous features are reported before the standardization was applied.

### Feature transformation and selection

Two groups of features correlated above the 0.75 threshold were identified, respectively the three ADAS scores (ADAS11, ADAS13, ADASQ4) and two of the RAVLT scores (RAVLT-F, RAVLT-PF). Such evidence equally resulted in all of the five training subsets. In all of the 5 subsets, only the first principal component of each group had an eigenvalue >= 1, and these were used to substitute the correlated features as predictors (ADAS-PC1, RAVLT-F-PC1).

### Code of the recruitment sites are those available in the ADNIMERGE file, following the coding convention used in the ADNI study

Across the five training subsamples used in the analyses, each feature selection procedure selected only partially overlapping subsets of relevant features, as reported in Table 3. Thus, the feature sets 2, 3, and 4 used in the analyses were in part different across the training subsamples used in the five repetitions of the analyses. This evidence further justifies our choice of creating several site-independent train and test subsamples instead of just a single training and test split, in order to provide a better and more stable estimate of the generalized performance of the algorithm.

**Table 3.**
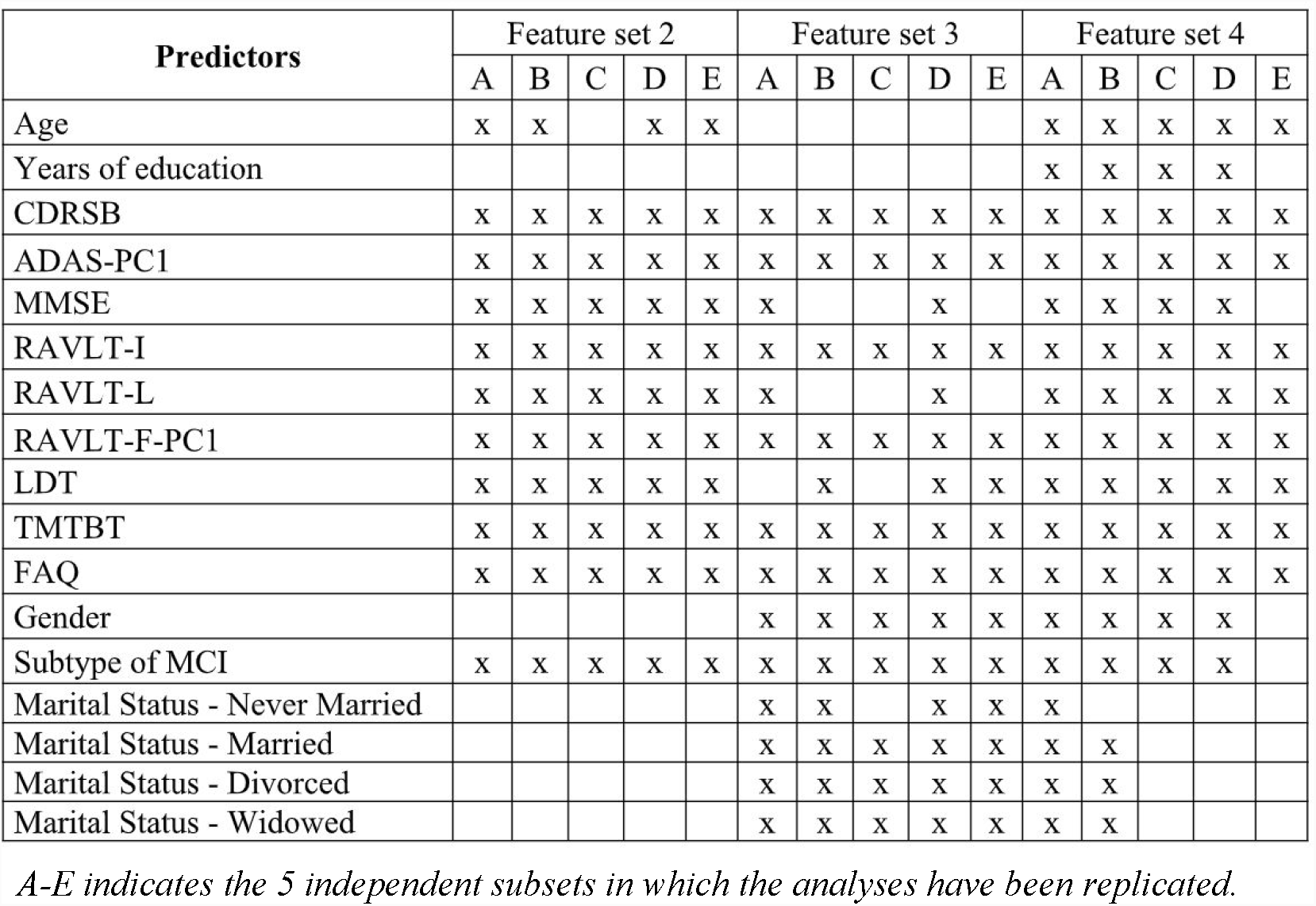
Feature sets 2, 3, and 4 in each of the five replications of the analyses.

### A-E indicates the 5 independent subsets in which the analyses have been replicated

Among the features, CDRSB, ADAS-PC1, RAVLT-I, RAVLT-F-PC1, TMTBT, and FAQ, were selected by all the three feature selection strategies in all of the five repetitions of the analyses, the subtype of MCI was discarded only once, LDT twice, RAVLT-L three times and MMSE four times. All the sociodemographic characteristics were all discarded at least 6 up to 11 times out of the 15 feature sets identified in the analyses.

### Performance of the predictive algorithm

The cross-validated AUROC results for each of the 52 models developed in each repetitions are reported in the supplementary data (Table S1), which ranged from a minimum value of 0.83 to a maximum value of 0.90 for the models developed with feature set 1, from 0.84 to 0.90 for the models developed with feature set 2, from 0.84 to 0.89 for the models developed with feature set 3, and from 0.83 to 0.90 for the models developed with feature set 4. These results indicate a narrow difference of performance among different feature sets, as well as among different replications and techniques, which included simple linear models such LR and NB as well as ensembling technique such as RF and GBM. The cross-validated AUROC of the weighted rank average ensembling strategy in each fold is also reported in Table S1, which ranged from a minimum of 0.86 to a maximum of 0.89.

When the test continuous prediction scores obtained with the ensembling approach were pooled, the whole sample test AUROC resulted 0.88 (95% bootstrap CI 0.85-0.91), which is plotted in Figure 1.

**Figure 1.**
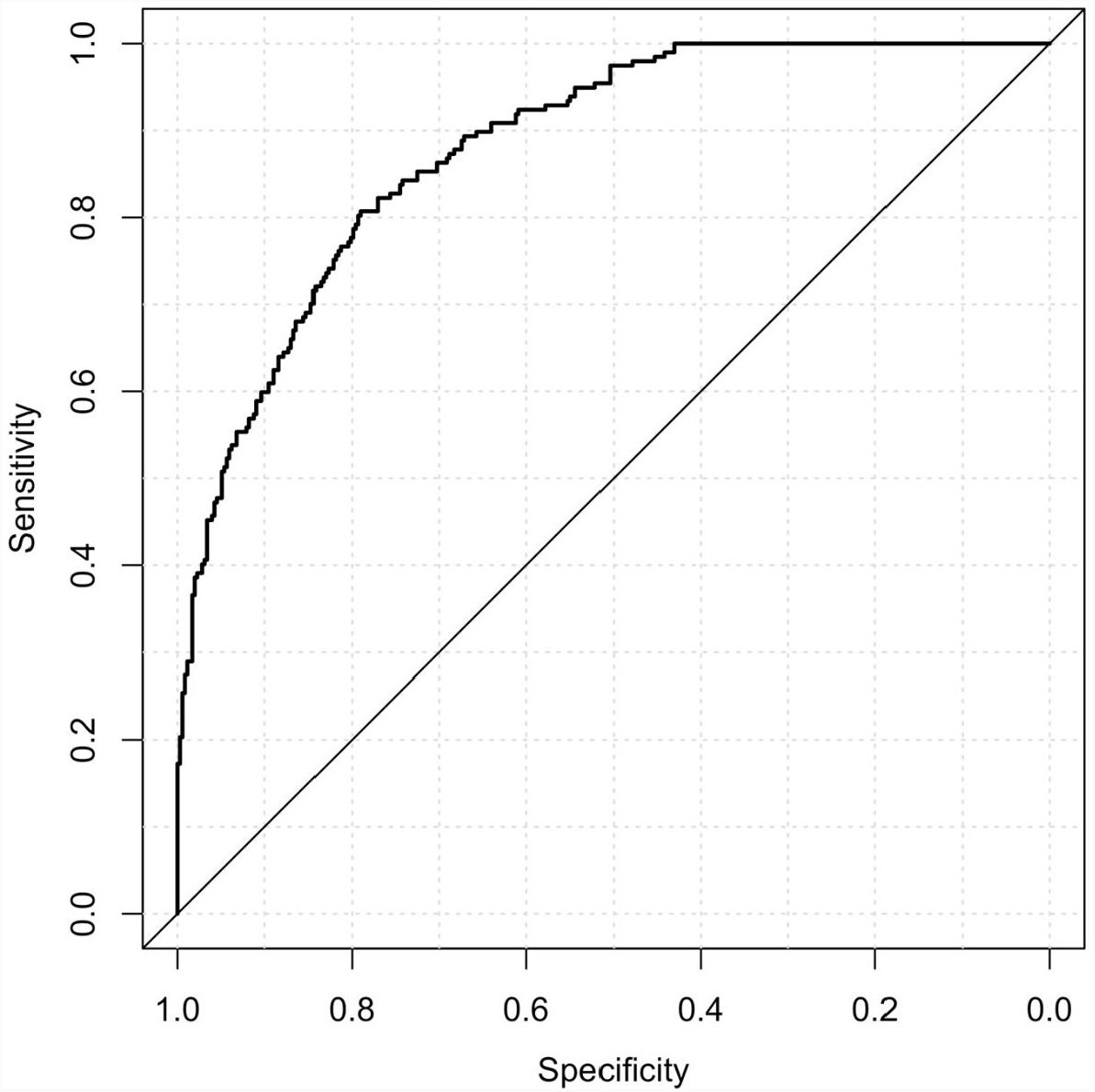
Area Under the Receiving Operating Curve of the pooled test predictions.

Considering the categorical predictions generated with the threshold that maximized the training balanced accuracy, results indicated a sensitivity/recall of 77.7%, a specificity of 79.9%, a positive predictive value/precision of 68.3%, a negative predictive value of 86.5%, a balanced accuracy of 0.79, and F1-score of 0.73. Results generated applying the other thresholds are reported in Table 4.

**Table 4.** Test performance of the algorithm.

| Aimed Sensitivity level | Sensitivity (Actual) | Specificity | Positive Predictive Value | Negative Predictive Value | Balanced Accuracy | F1-score |
| --- | --- | --- | --- | --- | --- | --- |
| Sensitivity of 1 | 100% | 40.2% | 48.3% | 100.0% | 0.701 | 0.651 |
| Sensitivity of .975 | 97.5% | 49.6% | 51.9% | 97.2% | 0.735 | 0.677 |
| Sensitivity of .95 | 94.9% | 53.0% | 53.0% | 94.9% | 0.739 | 0.680 |
| Sensitivity of .90 | 88.8% | 67.4% | 60.3% | 91.5% | 0.781 | 0.719 |
| Sensitivity of .85 | 84.3% | 73.1% | 63.6% | 89.3% | 0.787 | 0.725 |
| Sensitivity of .80 | 79.2% | 79.6% | 68.4% | 87.3% | 0.794 | 0.734 |
| Sensitivity of .75 | 71.6% | 84.1% | 71.6% | 84.1% | 0.779 | 0.716 |
| Best Balanced Accuracy | 77.7% | 79.9% | 68.3% | 86.5% | 0.788 | 0.727 |

*AUC = Area Under the Receiving Operating Curve*

All these results provide an estimate of the generalized performance of the algorithm when applied in new subjects which were not included in the sample used to develop the model and that have been evaluated in distinct recruiting sites.

### Importance of predictors

The AUROC of each of the various features obtained by pooling the results in the five test subsamples is reported in Table 5, ranked from the highest to the lowest AUROC, and in Figure 2, subdivided based on type of the features (i.e. sociodemographic, subtype of MCI, clinical, and neuropsychological tests). These represent an estimate of the generalized predictive performance achievable using each feature singularly.

**Table 5.** Individual test pooled AUROC of each feature.

|  | AUROC | 95% Bootstrap CI |  |
| --- | --- | --- | --- |
| ADAS-PC1 | 0.809 | 0.772 | 0.842 |
| RAVLT-I | 0.777 | 0.737 | 0.814 |
| FAQ | 0.777 | 0.733 | 0.816 |
| LDT | 0.770 | 0.726 | 0.808 |
| RAVLT-L | 0.707 | 0.661 | 0.750 |
| CDRSB | 0.697 | 0.648 | 0.740 |
| RAVLT-F-PC1 | 0.685 | 0.639 | 0.730 |
| MMSE | 0.678 | 0.631 | 0.723 |
| Subtype of MCI | 0.658 | 0.610 | 0.702 |
| TMTBT | 0.658 | 0.608 | 0.704 |
| Age | 0.564 | 0.511 | 0.614 |
| Years of education | 0.540 | 0.494 | 0.590 |
| Marital Status - Married | 0.506 | 0.452 | 0.547 |
| Marital Status - Divorced | 0.501 | 0.449 | 0.543 |
| Marital Status - Never Married | 0.488 | 0.439 | 0.537 |
| Marital Status - Widowed | 0.487 | 0.430 | 0.529 |
| Gender | 0.475 | 0.413 | 0.512 |

**Figure 2.**
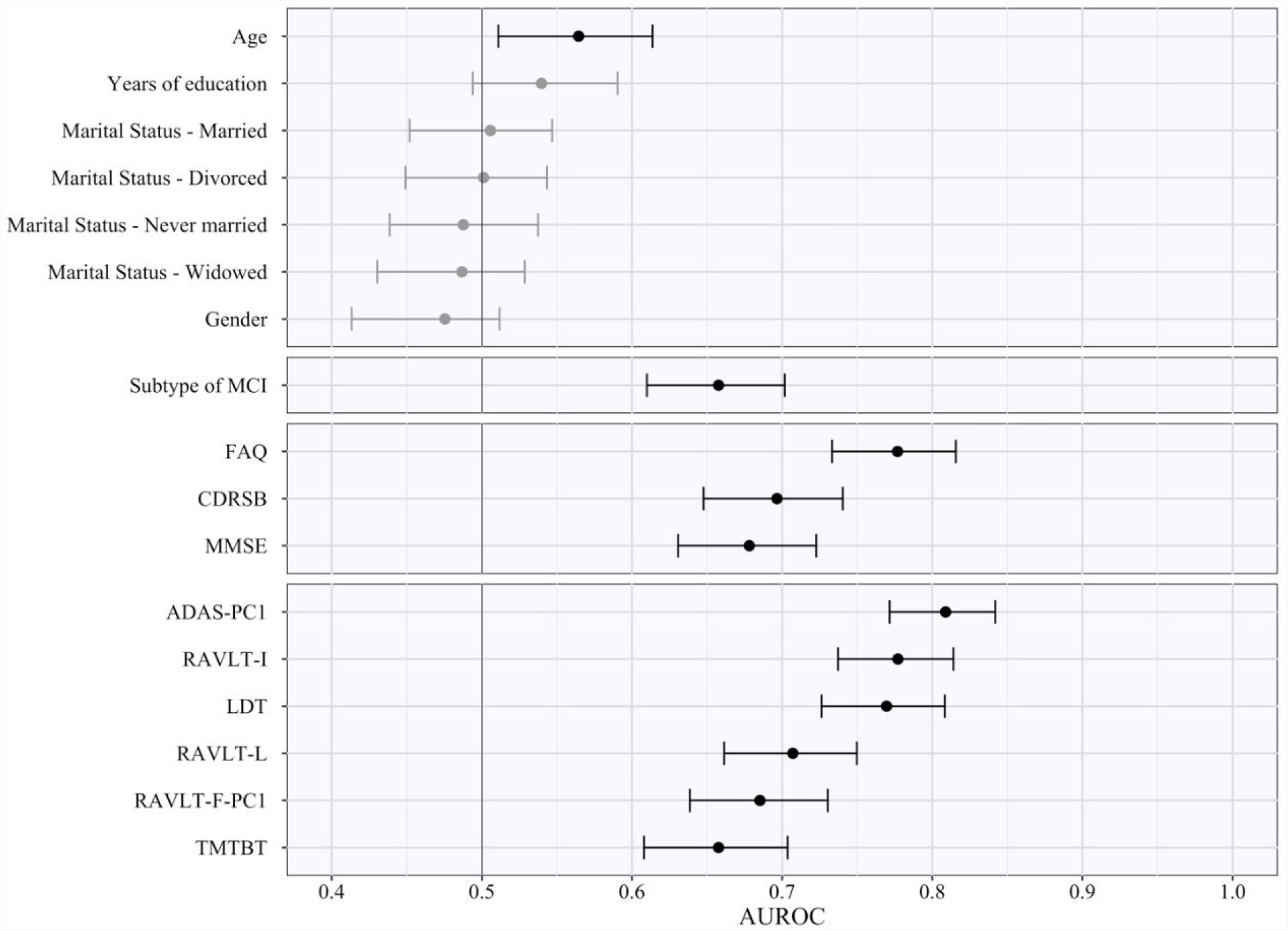
Area Under the Receiving Operating Curve of Individual Predictors. The figure indicates the pooled test AUROC and its 95% bootstrap CI when prediction is made considering each predictor singularly. Predictors are grouped according to conceptual domains, which in descending order are sociodemographic characteristics, subtype of MCI, clinical scale scores, and neuropsychological test scores. Non-significant AUCROC (i.e. the lower bound of the CI is lower than or equal to 0.5) are in grey, significant ones in black.

Sociodemographic characteristics resulted the least relevant, with age being the sole with a statistically significant AUROC (lower bound of the 95% bootstrap CI higher than 0.50) even if quite small in magnitude (AUROC_age_ = 0.57). Instead, both subtypes of MCI and CDRSB demonstrated a better predictive performance (AUROC_MCI_ = 0.66; AUROC_CDRSB_ = 0.70), and FAQ a high AUROC of 0.78. Among the neuropsychological test scores, some of them also proved to have a high predictive capability even when used as individual predictors. The ADAS-PC1 achieved an AUROC of 0.81, RAVLT-I of 0.78, and LDT of 0.77. All other neuropsychological test scores resulted with an inferior AUROC (minimum AUROC: AUROC_TMTBT_ = 0.66).

Of notice, the most relevant of the predictors, e.g. ADAS-PC1, resulted having a significantly lower test AUROC than the one demonstrated by the algorithm we developed (higher bound of the 95% bootstrap CI of ADAS-PC1 = 0.84 < lower bound of the 95% bootstrap CI of the algorithm = 0.85).

## Discussion

The aim of the current study was to develop a new machine-learning algorithm to allow a three-year prediction for conversion to AD in subjects diagnosed with MCI.

Considering an imminent necessity of being able to discriminate which MCI subjects will progress to Alzheimer’s from those who will not, as soon as in a few years the first effective treatments will be probably available [2], our algorithm has been designed to be used as a prognosis support tool for MCI patients, which is cost-effective and easily translatable to clinical practice. This would allow timely planning of early interventions for such individuals. Further, our algorithm can be employed as a tool during the recruitment of MCI subjects for clinical trials which aim to investigate innovative treatments of AD. The opportunity to recruit only subjects at true risk of future conversion to AD - who most likely show the earliest brain changes underlying AD pathology – will drastically reduce the costs to run such clinical trials and result in improved outcomes.

In contrast with many of the previous machine-learning approaches that have been previously presented, our algorithm aimed to achieve good predictive performance based only on a reduced set of sociodemographic characteristics, clinical information, and neuropsychological tests scores. It does not rely on information coming from procedures that are currently still expensive, invasive or not widespread available in many clinical settings, such as neuroimaging techniques, lumbar puncture, and genetic testing.

To develop the algorithm, we used data of 550 MCI subjects recruited in the ADNI study, and we applied a subdivision of the initial dataset in five mutually exclusive and collectively exhaustive, stratified test data subsets, in which the algorithm was iteratively tested after being trained in the remaining four subsets. This approach contributed to achieve an unbiased estimate of the predictive performance of our algorithm when it is applied to cases which were not used during the development phase of the algorithm. Moreover, we also ensured that the subsamples used to develop the algorithm contained subjects recruited from distinct sites than those from which subjects used for testing were recruited. This procedure was established in order to obtain results which represent an estimate of the expected performance, also when the algorithm is applied in distinct clinical centers. In addition, to the best of our knowledge, this is the first algorithm that was tested ensuring independence between the train and test sets regarding the sites where the subjects were recruited from.

Even using such a rigid testing protocol, the algorithm demonstrated a high predictive performance, showing a test AUROC of 0.88, and a test sensitivity and specificity of 77.7% and 79.9% respectively, when the algorithm was optimized during its development to achieve the best possible balanced accuracy. Of particular interest is the achievement of 40.2% specificity with a positive predictive value of 48.3% for 100% sensitivity, and a specificity and positive predictive value of 53% for 95% sensitivity. These results support the utility of our algorithm especially as a potential screening tool, i.e. an algorithm that can provide a marginal number of false negative predictions at the cost of a higher number of false positives. Thus, our algorithm would turn out to be particularly useful in case another more accurate, and especially more sensitive tool will become available, however which requires additional expensive or invasive-to-collect information. In such case, our algorithm can be used as a first step to significantly reduce the number of subjects which require examination using more precise, yet less easily applicable procedures at a later stage. Considering an expected conversion rate of 20%-40% from MCI to AD in three years, the expected percentage of subjects confidently predicted as non-converters would be estimated 32%-24% subsequently, leaving only the remaining 68%-76% of subjects with the necessity of further investigations.

Making a proper comparison of our algorithm with all others previously published is not a trivial task, especially considering the different and reduced level of independent validation most of these algorithms have undergone so far.

In some studies, algorithms which used as predictive information some type of functional brain imaging, such as PET and fMRI, and/or CSF investigations demonstrated particularly high cross-validated performance, with AUROCs close to 0.95 [17,18]. A recent study presented an algorithm based on regional information from a single amyloid PET scan which demonstrated a test performance of an AUROC of 0.91 and an unbalanced accuracy of 0.84 in the ADNI sample for a prediction of conversion in 2 years [42], thus showing a higher predictive performance than what was achieved by our algorithm.

In addition, some studies which used only structural MRI also demonstrated high cross-validated (i.e.[17,18]: AUROC = 0.932; balanced accuracy = 0.886) and nested cross-validated performance ([43]: sensitivity = 85%; specificity = 84.78%). Similarly, high cross-validated results were found by other studies who combined structural MRI with clinical and neuropsychological information (i.e. [7,10–18]): AUROC = 0.902; balanced accuracy = 80.5%) In addition, a recent study [44] presented a highly performing deep learning algorithm (AUROC = 0.925; accuracy = 86%; sensitivity = 87.5%; specificity = 85%) and, to the best of our knowledge, this is the only available study using structural MRI in which a a proper testing of the algorithm was performed.

Some particularly promising cross-validated results were also found in some studies which considered also APOE genotyping, together with EEG, ([45]: AUROC = 0.97; sensitivity = 96.7%; specificity = 86%) or blood biomarkers [7,10,16]: balanced accuracy = 92.5%). Thus, the use of brain imaging, CSF, and/or other biomarkers as predictive information may have, to some degree, resulted in a better predictive performance compared to our algorithm, which did not use any of these types of information.

While the results of the previous studies indicate that neuroimaging biomarkers hold great promise for predicting conversion to AD, the performance increase gained by including biomarker information is questioned and much debated [14,46,47]. Instead, neuropsychological measures of cognitive functioning are possibly equally excellent predictors of progression to dementia. For example, in a study by Fleisher and colleagues, common cognitive tests provide better predictive accuracy than imaging measures for predicting progression to AD in subject with moderate stages of amnestic MCI [47], and in another study by Clark and colleagues, models developed using only socio-demographic information, clinical information and neuropsychological test scores (focusing on verbal fluency scores) resulted in an AUROC score of 0.87 and a balanced accuracy of 0.84, while including brain imaging did not significantly improve this performance (AUROC = 0.81, accuracy = 0.83) [14].

Moreover, the cost of the standard procedure in the clinical process of diagnosing AD (which entails the clinical consultation, including the patient’s administrative admission, anamnesis, physical examination, neuropsychological testing, test evaluation and diagnosis conference & physician letter) is relatively low at an estimated 110 € (US$115) on average, while the use of additional advanced technical procedures, such as blood sampling, CT, MRI, PET & CSF procedures, which are required following deficits in neuropsychological test results and is dependent on the patient’s suspected diagnosis of MCI, AD or other dementia types (which is increasingly associated with higher frequencies of using cost-intensive imaging & CSF procedures), drives costs up to 649 € (US$676) in case of an AD diagnosis according to a study in a German memory clinic [48]. In this regards, the use of advanced technological procedures, rather than clinical consultation and neuropsychological testing, is driving costs in the diagnostic process and as such, will also increase the costs of predictive algorithms based on information of imaging, blood sampling or CSF procedures compared to those algorithms that rely only on sociodemographic, clinical, and neuropsychological predictive information, like the one we present in this study.

Additionally, our algorithm demonstrated similar predictive performance compared to other top-performing algorithms based only on sociodemographic, clinical, and neuropsychological predictive information. For example, in a first study by Clark and colleagues, they used only a simple cross-validation protocol to investigate the performance of their algorithm to make prediction of conversion at 1 year or more (AUROC = 0.88, balanced accuracy = 0.84) [13], while in another study they used a more sound nested cross-validation protocol to investigate the predictive performance of their algorithm at 4 years (AUROC = 0.87, balanced accuracy = 0.79) [14].

Altogether, our results originate from a proper testing protocol and represent a better unbiased estimate of the generalized performance of the algorithm. Only a very small number of machine learning algorithms for the prediction of conversion from MCI to AD were subjected to a proper testing protocol, rather than only a cross-validation protocol, which limits the soundness of the evidence of their predictive performance. As such, apart from [42,44], all the previously mentioned results may be optimistically biased estimates of the generalized performance of such algorithms as a proper testing protocol was not applied.

We previously presented another machine learning algorithm that performs a prediction of conversion to AD in MCI subjects [19,20]. However, the algorithm described here has distinct characteristics and can be considered at a more advanced stage of validation. First, the current algorithm does not require any neuroimaging information, while our previous method relied on a clinicians’ rating of the atrophy in three brain structures, evaluated by observing standardized images coming from a structural magnetic resonance. Structural magnetic resonance is widespread also in clinical settings nowadays, it is less expensive than other neuroimaging evaluation such as functional magnetic resonance and positron emission tomography, and the use of a clinician-administered visual scale allows to bypass the obstacles related to the non-automatic calibration of data coming from different magnetic resonance scanners. Nevertheless, the fact that our new algorithm does not necessitate any magnetic resonance evaluation makes its use even more easily translatable in practice, and less expensive. Moreover, even though our former algorithm showed higher cross-validated performance (AUROC = 0.91, sensitivity = 86.7% and specificity = 87.4% at the best balanced accuracy) [19]), a solid testing of its performance is still lacking and, at the moment, only a preliminary evidence via a transfer learning approach is available [20]. Instead, the protocol applied in the current study provides a better and sounder evaluation of the actual predictive performance of this new algorithm.

Beyond testing the algorithm’s predictive accuracy, we also aimed to provide a first indication of the importance of the variables used as predictors. The opportunity to provide an explanation of how the model works and performs its prediction is crucial to foster its application in clinical practice [49]. However, given the architectural complexity of the algorithm we developed, this is not a straightforward task. Several different approaches have been proposed, all of them providing different, and only a partial explanation of an algorithm’s functioning [50]. Thus, we decided to leave complex and more extensive investigations to a future study which will be fully dedicated to this goal. Instead, we simply investigated the predictive role of each predictor individually, which can evidence the amount of predictive information carried by each predictor. However, it does not allow to identify potential interactions among multiple predictors that could have been modeled by the algorithm and that can relevantly contribute to its high predictive performance.

In line with the evidence in our previous study [19], sociodemographic characteristics seem not to be particularly relevant in discriminating cAD and NC MCI subjects. Furthermore, in both studies, age was the sole of these characteristics showing a significant, even if very limited, predictive power. Also, sociodemographic characteristics resulted to be the most often discarded features by the feature selection strategies we applied in our study, once again suggesting their poor predictive relevance.

Instead, the clinical scale scores, the subtype of MCI, and the neuropsychological test scores resulted markedly predictive. Their test AUROC ranged from 0.658 to 0.809, and even the least predictive of them had a 95% CI higher than 0.6. The evidence of their predictive importance was expected. These features measure core elements of the progressive decline leading to a full manifestation of AD, such as the memory and other cognitive functions deterioration, and the consequent functional impairment.

Moreover, the first principal component of the three ADAS scores, which resulted in the most individually important predictor, demonstrated a test AUROC significantly lower than the one achieved by the entire algorithm. The results of our, as well as other previous studies, had already showed that machine learning algorithms can effectively be used to combine these individual pieces of information, providing a better identification of cAD among MCI subjects than what it would be possible using each of them singularly [13,14,19,20,46].

Our study has some limitations that should be taken into account and that will be addressed in the future stages of our research. First, even if we iteratively ensured that the subjects used for testing were always recruited in different sites than those used in the development of the algorithm, it is important to note that all the ADNI recruiting sites were located in the USA or Canada. Even if this can be considered an important step forward towards the demonstration of the generalized performance of the proposed algorithm, still these sites may not be completely representative of the entire population of centers in which the algorithm may aspire to be used. Our aim was to develop an algorithm that may be applied also beyond US and Canada centers only, and perhaps also clinical centers without any research inclinations. MCI subjects referring to these extended range of centers might have peculiar characteristics and the algorithm might show reduced predictive accuracy when applied to them. In order to at least partially address this potential bias, we plan to first test and then re-optimize our algorithm using further datasets coming from the several international replications of the North American ADNI (https://www.alz.org/research/for_researchers/partnerships/wwadni). In addition, inclusion and exclusion criteria may have excluded from ADNI, and in turn from our analyses, some MCI subjects with peculiar characteristics, e.g. MCI subjects with high level of depression or currently taking some of the medications that excluded for admission to the study. Once again, the algorithm might show reduced predictive accuracy when applied to them and further testing in a less selected sample should be performed before a safe use of the algorithm can be guaranteed with these peculiar MCI subjects.

Furthermore, our final algorithm is based on an ensemble of several lower-level machine learning algorithms, including some that use the entire initial set of predictors as feature set. Thus, all predictors currently remain necessary to be assessed, even if some of them may contribute poorly or even not at all to the prediction. Although the ensembling approach we used may have effectively prevented that such irrelevant predictors decreased the algorithm accuracy, a further reduction of the amount of information necessary to be assessed and used by the algorithm would permit to reduce the costs associated with its application.

Finally, our algorithm currently operates three-year predictions in subjects that already manifest MCI. As the new arriving treatments are expected to be the more effective the earlier they will be started, algorithms that can perform accurate predictions at even earlier stages of deterioration than MCI, and in a longer time frame, will be of particular relevance. A preliminary attempt has already been done in our previous study [19], employing also a sample of subjects with Pre-mild Cognitive Impairment [51], as well as in other previous studies which developed algorithm that aimed to make predictions for period longer than three years [10,14]. Future steps in our research will take into account this necessity, exploring the opportunity of making predictions at longer time periods and in earlier-stage subjects.

## Conclusions

We developed an algorithm to predict three-year conversion to AD in MCI subjects, based on a weighted rank average ensemble of several supervised machine learning algorithms. It demonstrated high predictive accuracy when tested via a sound train/test split protocol, exhibiting especially good predictive performance when the algorithm was optimized as a screening tool. Predictions are performed using only a restricted set of sociodemographic characteristics, clinical information, and neuropsychological test scores, which makes its application of easy translation into clinical practice, as well as useful in improving the recruitment of MCI subjects at true risk of conversion to AD in clinical trials. Further tests and optimizations will follow this study in order to provide additional evidence of its accuracy in generalized application, and to improve its cost-effectiveness.

## Abbreviations

AD: Alzheimer’s disease
ADNI: Alzheimer’s Disease Neuroimaging Initiative
ADAS: Alzheimer’s Disease Assessment Scale
ADAS11: Cognitive Subscale (11 items) Alzheimer’s Disease Assessment Scale
ADAS13: Cognitive Subscale (13 items) Alzheimer’s Disease Assessment Scale
ADASQ4: task 4 of the Cognitive Subscale (11 items) Alzheimer’s Disease Assessment Scale
AUROC: Area Under the Receiving Operating Curve
Bca: bias-corrected and accelerated
cAD: converters to probable Alzheimer’s disease
CDR: Clinical Dementia Rating Scale
CDRSB: Sum of Boxes score of the Clinical Dementia Rating Scale
CI: confidence interval
CSF: Cerebrospinal fluid
DIGIT: Digit Span Test score
EN: Elastic Net
FAQ: Functional Activities Questionnaire
GTB: Gradient Tree Boosting of Decision Trees
kNN: k-Nearest Neighbors algorithm
LDT: Logic Memory subtest of the of the Wechsler Memory Scale-Revised
LR: Logistic Regression
MLP1-Adam: Multi-Layer Perceptrons with one hidden layer and trained with adam algorithm
MLP1-Batch: Multi-Layer Perceptrons with one hidden layer and trained with full-batch gradient descent algorithm
MLP2-Adam: Multi-Layer Perceptrons with two hidden layers and trained with adam algorithm
MLP2-Batch: Multi-Layer Perceptrons with two hidden layers and trained with full-batch gradient descent algorithm
MCI: Mild cognitive impairment
MMSE: Mini-Mental State Examination
MRI: Magnetic resonance imaging
NB: Naive Bayes
NC: non-converters to Alzheimer’s disease
PET: Positron emission tomography
RAVLT: Rey Auditory Verbal Learning Test
RAVLT-F: Forgetting score of the Rey Auditory Verbal Learning Test
RAVLT-I: Immediate score of the Rey Auditory Verbal Learning Test
RAVLT-L: Learning score of the Rey Auditory Verbal Learning Test
RAVLT-PF: Percent forgetting score of the Rey Auditory Verbal Learning Test
RF: Random Forest
SVM-Linear: Support Vector Machine with linear kernel
SVM-RBF: Support Vector Machine with radial basis function kernel
SVM-Poly: and Support Vector Machine with polynomial kernel
TMTBT: Trail Making Test, version B.

## Acknowledgments

The authors would like to express their thanks to Lianne Ippel, Seun Onaopepo Adekunle, Alexander Malic, and Pedro Hernandez from the Institute of Data Science at Maastricht University for their valuable assistance, contributing their expertise to the experimental design of the analyses of this study. Data collection and sharing for this project was funded by the Alzheimer’s Disease Neuroimaging Initiative (ADNI) (National Institutes of Health Grant U01 AG024904) and DOD ADNI (Department of Defense award number W81XWH-12-2-0012). ADNI is funded by the National Institute on Aging, the National Institute of Biomedical Imaging and Bioengineering, and through generous contributions from the following: AbbVie, Alzheimer’s Association; Alzheimer’s Drug Discovery Foundation; Araclon Biotech; BioClinica, Inc.; Biogen; Bristol-Myers Squibb Company; CereSpir, Inc.; Cogstate; Eisai Inc.; Elan Pharmaceuticals, Inc.; Eli Lilly and Company; EuroImmun; F. Hoffmann-La Roche Ltd and its affiliated company Genentech, Inc.; Fujirebio; GE Healthcare; IXICO Ltd.; Janssen Alzheimer Immunotherapy Research & Development, LLC.; Johnson & Johnson Pharmaceutical Research & Development LLC.; Lumosity; Lundbeck; Merck & Co., Inc.; Meso Scale Diagnostics, LLC.; NeuroRx Research; Neurotrack Technologies; Novartis Pharmaceuticals Corporation; Pfizer Inc.; Piramal Imaging; Servier; Takeda Pharmaceutical Company; and Transition Therapeutics. The Canadian Institutes of Health Research is providing funds to support ADNI clinical sites in Canada. Private sector contributions are facilitated by the Foundation for the National Institutes of Health (www.fnih.org). The grantee organization is the Northern California Institute for Research and Education, and the study is coordinated by the Alzheimer’s Therapeutic Research Institute at the University of Southern California. ADNI data are disseminated by the Laboratory for NeuroImaging at the University of Southern California.

## Availability of data and material

Data used in the preparation of this article were obtained from the ADNI database (adni.loni.usc.edu), which is easily available for download from the Laboratory of Neuroimaging (LONI) website to the research public.

## Author Contributions

MG contributed to the design of the work, the design and execution of the analyses, interpretation of results, drafting of the paper and knowledge communication. NR contributed to the design of the work and the analyses, interpretation of results and drafting of the paper, as well as general project management, knowledge utilisation and communication, interactions with research professionals. DL contributed to the initial conception of the work and revising the manuscript. DC, KS, and GP contributed supervising the work and revising the manuscript. MD contributed to the design of the work and the analyses, interpretation of results, supervising the work and revising the manuscript. All authors read and approved the final manuscript.

## Declaration of Interest

The authors declare that they have no competing interests.

