## Supplementary Materials for "A Novel Ensemble-Based Machine Learning Algorithm To Predict The Conversion From Mild Cognitive Impairment To Alzheimer’s Disease Using Socio-demographic Characteristics, Clinical Information And Neuropsychological Measures"

**Table S1. Cross-validated AUROC of the algorithms developed in the five replications of the analyses.**

| Technique | Feature Set | Replications | | | | |
|---|---|---|---|---|---|---|
| | | A | B | C | D | E |
| k-Nearest Neighbor | 1 | 0.835 | 0.862 | 0.831 | 0.844 | 0.837 |
| Support Vector Machine, rbf kernel | | 0.867 | 0.893 | 0.871 | 0.886 | 0.873 |
| Support Vector Machine, polynomial kernel | | 0.865 | 0.892 | 0.870 | 0.883 | 0.871 |
| Support Vector Machine, linear kernel | | 0.867 | 0.893 | 0.871 | 0.886 | 0.873 |
| Random Forest | | 0.869 | 0.889 | 0.869 | 0.893 | 0.879 |
| Naïve Bayes | | 0.844 | 0.883 | 0.866 | 0.872 | 0.865 |
| Multi-layer Perceptron, 2 hidden layers and batch learning | | 0.860 | 0.885 | 0.872 | 0.883 | 0.873 |
| Multi-layer Perceptron, 2 hidden layers and adam learning | | 0.865 | 0.895 | 0.879 | 0.898 | 0.875 |
| Multi-layer Perceptron, 1 hidden layer and batch learning | | 0.862 | 0.888 | 0.876 | 0.888 | 0.870 |
| Multi-layer Perceptron, 1 hidden layer and adam learning | | 0.867 | 0.896 | 0.880 | 0.902 | 0.876 |
| Logistic Regression | | 0.865 | 0.886 | 0.869 | 0.886 | 0.870 |
| Gradient Tree Boosting | | 0.868 | 0.894 | 0.872 | 0.892 | 0.884 |
| Elastic Net | | 0.855 | 0.885 | 0.867 | 0.876 | 0.872 |
| k-Nearest Neighbor | 2 | 0.851 | 0.879 | 0.863 | 0.880 | 0.868 |
| Support Vector Machine, rbf kernel | | 0.872 | 0.896 | 0.878 | 0.888 | 0.878 |
| Support Vector Machine, polynomial kernel | | 0.873 | 0.898 | 0.882 | 0.889 | 0.886 |
| Support Vector Machine, linear kernel | | 0.872 | 0.896 | 0.879 | 0.888 | 0.878 |
| Random Forest | | 0.878 | 0.894 | 0.871 | 0.892 | 0.877 |
| Naïve Bayes | | 0.843 | 0.876 | 0.856 | 0.854 | 0.863 |
| Multi-layer Perceptron, 2 hidden layers and batch learning | | 0.863 | 0.902 | 0.877 | 0.888 | 0.879 |

| | | | | | | |
|---|---|---|---|---|---|---|
| Multi-layer Perceptron, 2 hidden layers and adam learning | | 0.867 | 0.895 | 0.864 | 0.881 | 0.869 |
| Multi-layer Perceptron, 1 hidden layer and batch learning | | 0.869 | 0.899 | 0.877 | 0.892 | 0.880 |
| Multi-layer Perceptron, 1 hidden layer and adam learning | | 0.862 | 0.896 | 0.876 | 0.889 | 0.879 |
| Logistic Regression | | 0.870 | 0.893 | 0.872 | 0.884 | 0.869 |
| Gradient Tree Boosting | | 0.876 | 0.897 | 0.877 | 0.894 | 0.887 |
| Elastic Net | | 0.871 | 0.898 | 0.876 | 0.887 | 0.877 |
| k-Nearest Neighbor | | 0.838 | 0.861 | 0.851 | 0.852 | 0.847 |
| Support Vector Machine, rbf kernel | | 0.866 | 0.894 | 0.875 | 0.886 | 0.872 |
| Support Vector Machine, polynomial kernel | | 0.866 | 0.893 | 0.876 | 0.885 | 0.871 |
| Support Vector Machine, linear kernel | | 0.866 | 0.894 | 0.874 | 0.886 | 0.872 |
| Random Forest | | 0.869 | 0.890 | 0.874 | 0.890 | 0.879 |
| Naïve Bayes | | 0.863 | 0.892 | 0.867 | 0.873 | 0.869 |
| Multi-layer Perceptron, 2 hidden layers and batch learning | 3 | 0.862 | 0.892 | 0.874 | 0.879 | 0.877 |
| Multi-layer Perceptron, 2 hidden layers and adam learning | | 0.866 | 0.894 | 0.875 | 0.890 | 0.876 |
| Multi-layer Perceptron, 1 hidden layer and batch learning | | 0.865 | 0.891 | 0.874 | 0.888 | 0.877 |
| Multi-layer Perceptron, 1 hidden layer and adam learning | | 0.864 | 0.893 | 0.868 | 0.894 | 0.875 |
| Logistic Regression | | 0.865 | 0.891 | 0.868 | 0.884 | 0.867 |
| Gradient Tree Boosting | | 0.868 | 0.893 | 0.875 | 0.892 | 0.883 |
| Elastic Net | | 0.861 | 0.889 | 0.873 | 0.879 | 0.869 |
| k-Nearest Neighbor | | 0.835 | 0.865 | 0.834 | 0.852 | 0.848 |
| Support Vector Machine, rbf kernel | | 0.867 | 0.892 | 0.870 | 0.886 | 0.872 |
| Support Vector Machine, polynomial kernel | | 0.865 | 0.892 | 0.869 | 0.883 | 0.871 |
| Support Vector Machine, linear kernel | | 0.867 | 0.892 | 0.870 | 0.886 | 0.872 |
| Random Forest | | 0.869 | 0.890 | 0.868 | 0.892 | 0.879 |
| Naïve Bayes | | 0.844 | 0.873 | 0.844 | 0.854 | 0.858 |
| Multi-layer Perceptron, 2 hidden layers and batch learning | 4 | 0.860 | 0.890 | 0.865 | 0.894 | 0.877 |

| | | | | | | |
|---|---|---|---|---|---|---|
| Multi-layer Perceptron, 2 hidden layers and adam learning | | 0.865 | 0.894 | 0.876 | 0.892 | 0.876 |
| Multi-layer Perceptron, 1 hidden layer and batch learning | | 0.862 | 0.892 | 0.871 | 0.882 | 0.870 |
| Multi-layer Perceptron, 1 hidden layer and adam learning | | 0.867 | 0.895 | 0.879 | 0.898 | 0.877 |
| Logistic Regression | | 0.865 | 0.886 | 0.864 | 0.882 | 0.862 |
| Gradient Tree Boosting | | 0.868 | 0.893 | 0.874 | 0.892 | 0.882 |
| Elastic Net | | 0.858 | 0.885 | 0.868 | 0.876 | 0.865 |
| Weighted Rank Average | | 0.858 | 0.891 | 0.876 | 0.885 | 0.882 |

Rbf = radial-basis function kernel.
